# Spatially distinct inputs modulate the amount of active Mitotic-phase GAP to locally restrict RhoA signaling for successful cell division

**DOI:** 10.1101/2023.08.08.552464

**Authors:** Friederike Wolff, Swathi Srinivasan, Marie Christin Nöcker, Sriyash Mangal, Tamara Mikeladze-Dvali, Esther Zanin

## Abstract

At the end of mitosis a contractile ring consisting of filamentous actin (F-actin) assembles at the cell equator and ring constriction equally partitions the cellular content. Inhibitory and stimulatory signaling cascades spatially limit RhoA activity to a narrow central zone to promote F-actin polymerization only at the cell equator. While the stimulatory signal is well-characterized, the mechanisms by which inhibitory signals restrict active RhoA to a narrow equatorial zone and prevent improper RhoA activity at the cell poles are not known. Here, we identify two regulatory inputs that control the activation and cortical targeting of the Mitotic-phase GTPase activating protein (MP-GAP) for RhoA which limits RhoA activity both at the cell poles and at the cell equator. We show that at cell poles, MP-GAP is a direct phosphorylation target of spindle-pole associated Aurora A kinase. We identify three Aurora A target residues in a region of MP-GAP that binds directly to its catalytic GAP domain suggesting the Aurora A phosphorylation releases MP-GAP autoinhibition. We show that phosphorylation of these sites is required to keep RhoA activity low at the cell poles. At the cell equator, F-actin facilitates MP-GAP enrichment, which opposes RhoA GEF activity and thereby accelerates RhoA flux through the GTPase cycle for the formation of a narrow equatorial zone of active RhoA. Thus, the amount of active MP-GAP is modulated by two distinct regulatory inputs that function in spatially restricted locations: Aurora A phosphorylation relieves MP-GAP autoinhibition to limit RhoA activity at the cell poles, while F-actin polymerization promotes MP-GAP targeting to prevent distribution of active RhoA in a broad zone at the cell equator. By determining the mechanism of spatially confining RhoA activity at the equator and the cell poles during cytokinesis, our work has broad implications to how Rho activity zones are formed and maintained during cytokinesis and how defects in their formation impact animal development and disease.

## Introduction

During the last step of cell division, called cytokinesis, the content of the mother cell is segregated between the two daughter cells. A contractile ring consisting of filamentous actin (F-actin) and non-muscle myosin II assembles at the cell equator during anaphase and ring constriction forms the two daughter cells (Basant and Glotzer, 2018; Mishima, 2016; Pollard and O’Shaughnessy, 2019). Contractile ring formation is induced by the small GTPase RhoA, which is active in a narrow zone at the cell equator. Opposing stimulatory and inhibitory signals from spindle-associated proteins restrict RhoA activity to a narrow equatorial zone and thereby tightly link chromosome segregation with cleavage plane specification. The well characterized stimulatory signal derives from the spindle midzone, which consists of overlapping antiparallel microtubule bundles forming between the segregating chromosomes. The stimulatory signal releases autoinhibition of the RhoA GEF Ect2 and thereby promotes RhoA activation at the cell equator (Burkard et al., 2009; Chen et al., 2019; Dechant and Glotzer, 2003; Gomez-Cavazos et al., 2020; Prokopenko et al., 1999; Schneid et al., 2021; Somers and Saint, 2003; Tatsumoto et al., 1999; Wolfe et al., 2009; Yüce et al., 2005). Whereas the molecular mechanisms of the stimulatory signal are known, how the inhibitory signals restrict RhoA activity to a narrow zone at the cell equator and limit RhoA activity at the cell poles are less understood.

It has been demonstrated decades ago that the spindle poles can induce furrowing without being connected by a spindle in a seminal paper by Rappaport (Rappaport, 1961). Based on those observations it was postulated that the spindle poles are the source of an inhibitory signal that induces relaxation of the cell poles (White and Borisy, 1983; Wolpert, 1960). Since then, numerous studies in different organisms have confirmed the presence of a spindle pole-based inhibitory signal. For example, ablation of the centrosome or selective removal of astral microtubules broadens the zone of active RhoA at the cell equator (Bement et al., 2005; Foe and von Dassow, 2008; Murthy and Wadsworth, 2008; van Oostende Triplet et al., 2014; von Dassow et al., 2009; Zanin et al., 2013). Shifting the position of the spindle poles by genetic or mechanical manipulation to one site causes an accumulation of contractile ring proteins at the opposite site (Chen et al., 2008; Werner et al., 2007). Similarly, altering spindle pole position by laser microsurgery causes the furrow site to shift in position (Bringmann and Hyman, 2005; von Dassow et al., 2009). In contrast, increasing the number of astral microtubules by depleting the microtubules depolymerizing kinesin MCAK narrows the RhoA zone (van Oostende Triplet et al., 2014; Zanin et al., 2013). The reduction in cortical contractility at the cell poles was proposed to induce equatorial directed flows which cause equatorial compression of the cortex and consequently a positive feedback loop that mediates ring ingression (Khaliullin et al., 2018; Murthy and Wadsworth, 2005; Reymann et al., 2016; Zhou and Wang, 2008).

We found that the Aurora A kinase activator, TPXL-1 (TPX2 in humans), promotes clearing of contractile ring components from the cell poles in the *C. elegans* one-cell embryo (Mangal et al., 2018). TPXL-1/TPX2 binds astral microtubules where it activates Aurora A via its N-terminus (Bayliss et al., 2003; Kufer et al., 2002). Aurora A is a major mitotic kinase involved in mitotic entry, centrosome maturation and spindle assembly (Tavernier et al., 2021). Aurora A localizes to astral microtubules and centrosomes and is active in a gradient around monopolar spindles (Kufer et al., 2002; Portier et al., 2007; Stenoien et al., 2003; Sugimoto et al., 2002; Ye et al., 2015). Thus, active Aurora A could diffuse from the spindle poles to the overlaying cell cortex where it inhibits RhoA signaling. In the *C. elegans* zygote, a recent report proposes that Aurora A phosphorylation of Ect2 in its membrane binding PH domain removes Ect2 from the posterior pole during anaphase causing an accumulation of Ect2 at the opposite anterior membrane (Longhini and Glotzer, 2022). How Ect2 activity is shut off at the anterior membrane to avoid polar RhoA activity remains unclear. Additionally, astral microtubules serve as tracks for dynein dependent myosin II removal from the cell poles in *C. elegans* (Chapa-y-Lazo et al., 2020).

Besides spindle pole derived signals, other mechanisms were also shown to inhibit polar contractility. The kinetochore localized phosphatase PP1 inactivates ezrin/radixin/moesin at the cell poles and thereby softens the polar cortex (Rodrigues et al., 2015). Another chromatin localized signal comprises the GTPase Ran, which blocks binding of contractile ring components to the polar regions (Beaudet et al., 2017; Beaudet et al., 2020; Kiyomitsu and Cheeseman, 2013; Ozugergin et al., 2021; Ozugergin et al., 2022).

The key negative regulator of RhoA activity during mitosis, which increases the low intrinsic GTP hydrolysis rate of RhoA, is MP-GAP (also named ArhGAP11a). MP-GAP is conserved from *C. elegans* to human and localizes to the cell periphery in metaphase and enriches at the cell equator during anaphase (Michaud et al., 2022; Zanin et al., 2013). *C. elegans* has two MP-GAP homologues called RGA-3 and RGA-4 (RGA-3/4) which limit the accumulation of contractile ring proteins to the cell cortex during mitosis and thereby redundantly contribute to successful cytokinesis (Bell et al., 2020; Schmutz et al., 2007; Schonegg et al., 2007; Zanin et al., 2013). RGA-3/4 localization to the cell periphery requires an intact F-actin cortex and RGA-3/4 terminate RhoA activity pulses during polarization in *C. elegans* and in frog and starfish oocytes during mitosis (Michaud et al., 2022; Michaux et al., 2018). In human cells, MP-GAP limits cortical contractility and in case the spindle pole-based inhibitory signal is reduced, MP-GAP restricts RhoA zone dimension during cytokinesis (Zanin et al., 2013).

Thus, the molecular mechanisms of how inhibitory signals spatially confine RhoA activity during mitosis remain sparse. Here we reveal that Aurora A kinase is part of a spindle-pole localized inhibitory signal in human cells and blocks RhoA signaling at the cell poles. Using microscopy and biochemistry-based assays we demonstrate that Aurora A limits RhoA signaling at the cell poles by phosphorylating MP-GAP in a region adjacent to its catalytic GAP domain. Binding studies show that this region of MP-GAP binds to its catalytic GAP domain suggesting that MP-GAP adopts an autoinhibitory conformation, which is released by Aurora A kinase phosphorylation. Furthermore, our data also demonstrate that MP-GAP not only functions at the cell poles but also at the cell equator. MP-GAP is targeted to the cell equator by F-actin, and due to high activation of RhoA by Ect2 in this region, equatorial MP-GAP accelerates RhoA flux through the GTPase cycle and thereby narrows the width of the zone of active RhoA. In summary, we discover that MP-GAP is regulated by two spatially distinct regulatory inputs: spindle-pole dependent activation by Aurora A and equatorial recruitment by F-actin. Together these two inputs spatially fine tune the amount of active MP-GAP and thereby contribute to the formation of a narrow RhoA zone at the cell equator and low RhoA activity at the cell poles.

## Results

### Aurora A kinase restricts anillin accumulation at the cell poles and to a narrow equatorial zone during anaphase in human cells

The Aurora A kinase activator TXPL-1 promotes clearing of contractile ring components from the cell poles in *C. elegans* (Mangal et al., 2018). To test whether the function of Aurora A is conserved in human cells (Fig. 1A), we treated HCT-116 cells, a colon cancer cell line expressing endogenously NeonGreen-tagged anillin (Husser et al., 2022) with the highly specific Aurora A inhibitor MK-5108 (de Groot et al., 2015). Since anillin binds active RhoA it represents an excellent readout for active RhoA localization (Budnar et al., 2019; Piekny and Glotzer, 2008; Sun et al., 2015). To exclude that the observed phenotype arises from the inhibition of highly similar Aurora B kinase by MK-5108, we also inhibited Aurora B for comparison. For Aurora B inhibition we chose AZD1152-HQPA (AZD), which was shown to be highly selective for Aurora B (de Groot et al., 2015). MK-5108 and AZD were added during live-cell imaging before anaphase onset and as expected Aurora A but not Aurora B inhibition reduced spindle midzone length during anaphase confirming previous findings (Fig. S1A, (Reboutier et al., 2013)). Furthermore, cytokinesis failure was only observed after AZD but not after MK-5108 treatment (Fig. S1B, (Carmena et al., 2012)). Since MK-5108 and AZD treatment resulted in distinct and previously published phenotypes, we are confident that our treatment conditions are highly specific for each kinase and thus we continued to analyze anillin localization. In control anaphase cells treated with DMSO, anillin is strongly enriched in a narrow equatorial zone and weakly present at the cell poles (Fig. 1B, C). After Aurora A, but not Aurora B, inhibition the equatorial anillin zone appeared broader and more anillin accumulated at the cell poles. To quantitatively compare anillin distribution, we measured anillin fluorescence intensity around the cell periphery (Fig. 1B) and confirmed that polar anillin levels were increased and the equatorial anillin zone was broader after Aurora A, but not Aurora B inhibition (Fig. 1C). Equatorial levels of anillin were not increased after Aurora A or Aurora B inhibition (Fig. 1C). A broader spindle midzone results in a broader equatorial anillin zone (Hu et al., 2011), however after Aurora A inhibition the spindle midzone length was shorter and thus cannot explain the broadened anillin zone. To ensure that Aurora A inhibition does not indirectly alter anillin localization by reducing the number of astral microtubules, we stained anaphase cells for α-tubulin. In MK-5108-treated HCT-116 cells the spindle midzone was disorganized (Reboutier et al., 2013) but the astral microtubules were still present and did not exhibit any difference to astral microtubules in control cells (Fig. S1C). Aurora A is present on spindle microtubules and centrosomes already during metaphase (Kufer et al., 2002; Sugimoto et al., 2002) and therefore we asked whether Aurora A inhibition during metaphase also increases cortical anillin levels. HCT-116 cells expressing NG-tagged anillin were treated during live-cell imaging with each inhibitor during prometaphase and cortical anillin levels at metaphase were analyzed. Indeed, cortical anillin levels were also increased in metaphase cells after Aurora A, but not Aurora B inhibition (Fig. S1D).

**Figure 1.**
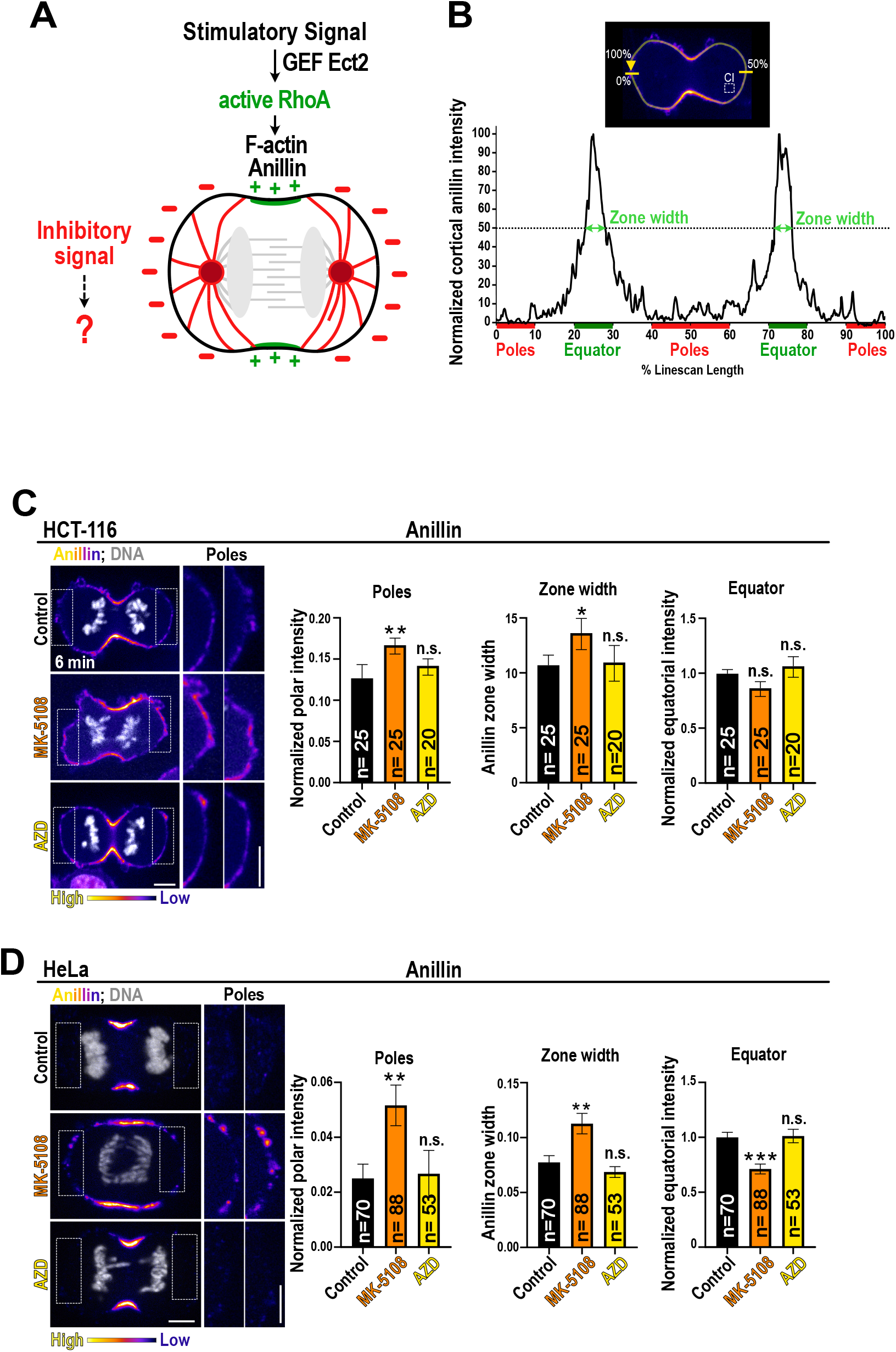
Aurora A kinase limits anillin accumulation at the cell poles and to a narrow equatorial zone in human cells. A) Scheme of the stimulatory and inhibitory spindle-derived signals that spatially pattern RhoA activity during cytokinesis. B) A linescan was drawn around the cell periphery, starting at one cell pole to quantify anillin, fluorescence intensity along the cortex. Anillin cytoplasmic intensity (CI) was measured in a box in the cytoplasm. The normalized anillin fluorescence intensity from 0%–100% line scan length is shown for a control HCT-116 cell. The polar anillin intensity was the mean intensity from 0%-10%, 40-60% and 90-100% line scan length, the equatorial intensity the mean from 40-50% and 70-80%, and the anillin zone width was determined at the half maximum fluorescence intensity. C) Shown are time-lapse fluorescent images of HCT-116 cells expressing NG-Anillin (pseudocolour) labeled for DNA (SiR-DNA, grey) 6 min after anaphase onset. HCT-116 cells were treated with MK-5108, AZD, or DMSO (control). Insets show a magnification of the poles. Plotted is the mean normalized fluorescence intensity of anillin at the cell poles, the equator and the width of the equatorial anillin zone in % linescan length 6 min after anaphase onset. D) Immunofluorescent images of anaphase HeLa cells labeled for endogenous anillin (pseudocolor) and DNA (grey) treated with MK-5108, AZD, or DMSO (control). Insets show a magnification of the poles. Intensity scalings are increased for the magnifications to better visualize polar anillin accumulation. Plotted is the mean normalized fluorescence intensity of anillin at the cell poles, the cell equator and the width of the equatorial anillin zone. For all n = number of cells, and ≥3 independent experiments were performed for each condition. Error bars are SEM, and p values were calculated with the Kruskal-Wallis test. Scale bars are 5 µm.

Finally, we investigated whether Aurora A limits anillin accumulation during mitosis also in human cervical cancer HeLa cells. Like in HCT-116 cells, Aurora A inhibition resulted in a shorter spindle midzone length and Aurora B inhibition caused cytokinesis failure (Fig. S1E, F). Immunostaining of endogenous anillin revealed that after Aurora A inhibition polar anillin levels were increased, the width of the anillin zone was widened, and equatorial anillin levels were decreased in comparison to control anaphase cells (Fig. 1D). Similar to HCT-116 cells, cortical anillin levels were also elevated at metaphase after Aurora A inhibition (Fig. S1G). Although Aurora B inhibited cells frequently failed in cytokinesis, they did not show any alteration in anillin localization during mitosis (Fig. 1D, S1G).

In summary, Aurora A limits the cortical anillin accumulation at the cell poles and maintains anillin in a narrow equatorial zone during anaphase, suggesting that the inhibitory role of Aurora A at the cell poles is conserved in human cells.

### Aurora A kinase phosphorylates S285, S304, S345 of MP-GAP

Next, we aimed to identify the phosphorylation target of Aurora A kinase that limits RhoA signaling at the cell poles. We hypothesized that Aurora A promotes RhoA inactivation by activating the negative RhoA regulator MP-GAP (Fig. 2A). To test if Aurora A phosphorylates MP-GAP, we split MP-GAP into three fragments comprising the catalytic GAP domain (GST-GAP: 1-253 AA), the F1 (GST-F1: 254-640 AA), or F2 (GST-F2: 640-1021 AA, Fig. 2B) region.

**Figure 2.**
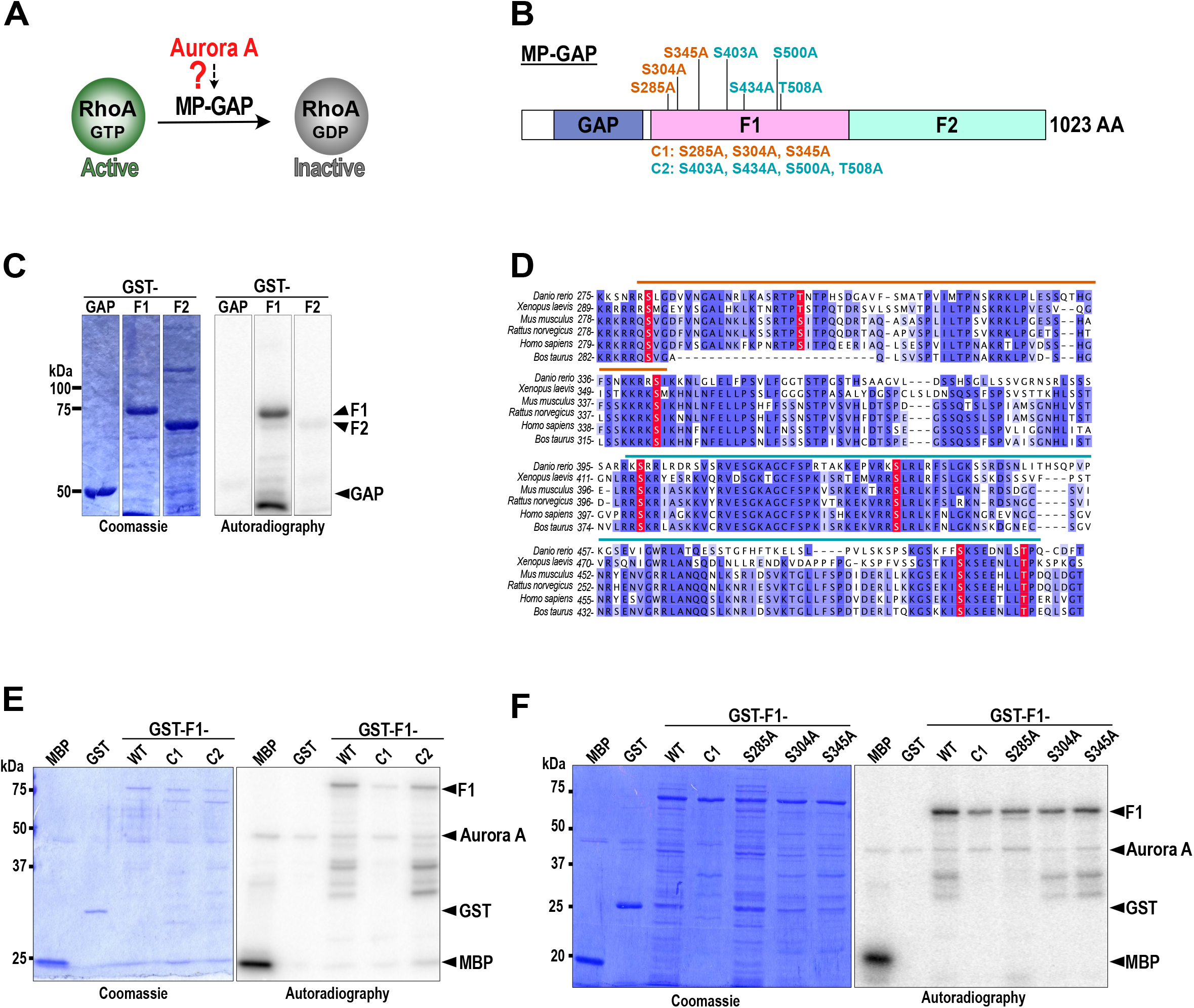
Aurora A kinase phosphorylates S285, S304 and S345 of MP-GAP. A) Aurora A kinase could facilitate RhoA inactivation by activating the RhoA GAP MP-GAP. B) Schematic of MP-GAP domain organization highlighting the three fragments and putative phosphosites tested in the kinase assays. C) Coomassie gel and autoradiography images of Aurora A kinase assays with GST-tagged GAP domain and F1 and F2 region of MP-GAP. Different lanes of the same gel are shown. D) Sequence alignment of the F1 region of MP-GAP. The conserved putative Aurora A phosphosites are highlighted in red. E, F) Coomassie gel and autoradiography images of Aurora A kinase assays with the myelin basic protein (MBP, positive control), GST (negative control), GST-tagged F1-WT, F1-C1 (S285A, S304A, S345A) and F1-C2 (S403A, S434A, S500A, T508A) (E) or F1-S285A, F1-S304A, and F1-S345A (F) of MP-GAP.

Each fragment was incubated with Aurora A kinase and their phosphorylation was tested by *in vitro* kinase assays. We found that GST-F1, but not GST-GAP or GST-F2 was phosphorylated by Aurora A kinase (Fig. 2C). To narrow down which amino acids are phosphorylated, we used the group-based prediction system (GPS 5.0) program and identified seven conserved Aurora kinase consensus phosphorylation sites in F1 region (Fig. 2D). To test which sites are phosphorylated we grouped them into two clusters and generated phosphodeficient cluster 1 (C1: S285A, S304A, S345A) and cluster 2 (C2: S403A, S434A, S500A, T508A) GST-F1 variants (Fig. 2B). After incubation of the GST-F1-C1, but not GST-F1-C2 mutant with Aurora A, we observed a strong reduction in the amount of phosphorylation (Fig. 2E). We finally tested whether any single-site-mutation to alanine in C1 reduced Aurora A kinase phosphorylation. Indeed, we observed that each single-site-mutation in C1 (S285A, S304A, or S345A) reduced Aurora A kinase phosphorylation (Fig. 2F), suggesting that each site is phosphorylated. In sum, Aurora A kinase phosphorylates S285, S304, and S345 in the F1 region of MP-GAP *in vitro*.

### MP-GAP phosphorylation at S285, S304, and S345 limits anillin accumulation at the cell poles and to a narrow equatorial zone

After Aurora A inhibition cortical anillin levels were increased at the cell poles and its equatorial zone was broadened. Since Aurora A phosphorylates MP-GAP *in vitro*, we asked whether MP-GAP depletion mimics the Aurora A inhibition phenotype. After MP-GAP depletion cortical anillin levels were increased during metaphase (Fig. 3A). Similarly, during anaphase the cortical levels of anillin at the cell poles were also elevated and the equatorial zone width was increased, however the equatorial anillin levels were unchanged (Fig. 3B, C). To test if Aurora A phosphorylation of S285, S304, and S345 is required for MP-GAP function, we made use of a genetic replacement system, which consists of a stably integrated RNAi-resistant GFP-tagged MP-GAP wild-type (MP-GAP^WT^) transgene in HeLa cells (Fig. S2A, B, (Zanin et al., 2013)). Additionally, we generated a cell line expressing a full length phosphodeficient MP-GAP mutant transgene in which S285, S304, and S345 of MP-GAP were mutated to alanine (MP-GAP^S3A^, Fig. S2A). By performing western blotting, we confirmed that after MP-GAP RNAi, endogenous MP-GAP was strongly reduced and MP-GAP^WT^ and MP-GAP^S3A^ were expressed at similar levels as endogenous MP-GAP (Fig. S2B). Finally, we analyzed cortical anillin localization during mitosis: we observed that after MP-GAP RNAi expression of MP-GAP^WT^, but not MP-GAP^S3A^, reduced cortical anillin levels in metaphase (Fig. 3A). Similarly, during anaphase MP-GAP^WT^ but not MP-GAP^S3A^ expressing cells treated with MP-GAP RNAi had lower cortical anillin levels at the cell poles and a control-like anillin zone at the cell equator (Fig. 3B, C).

**Figure 3.**
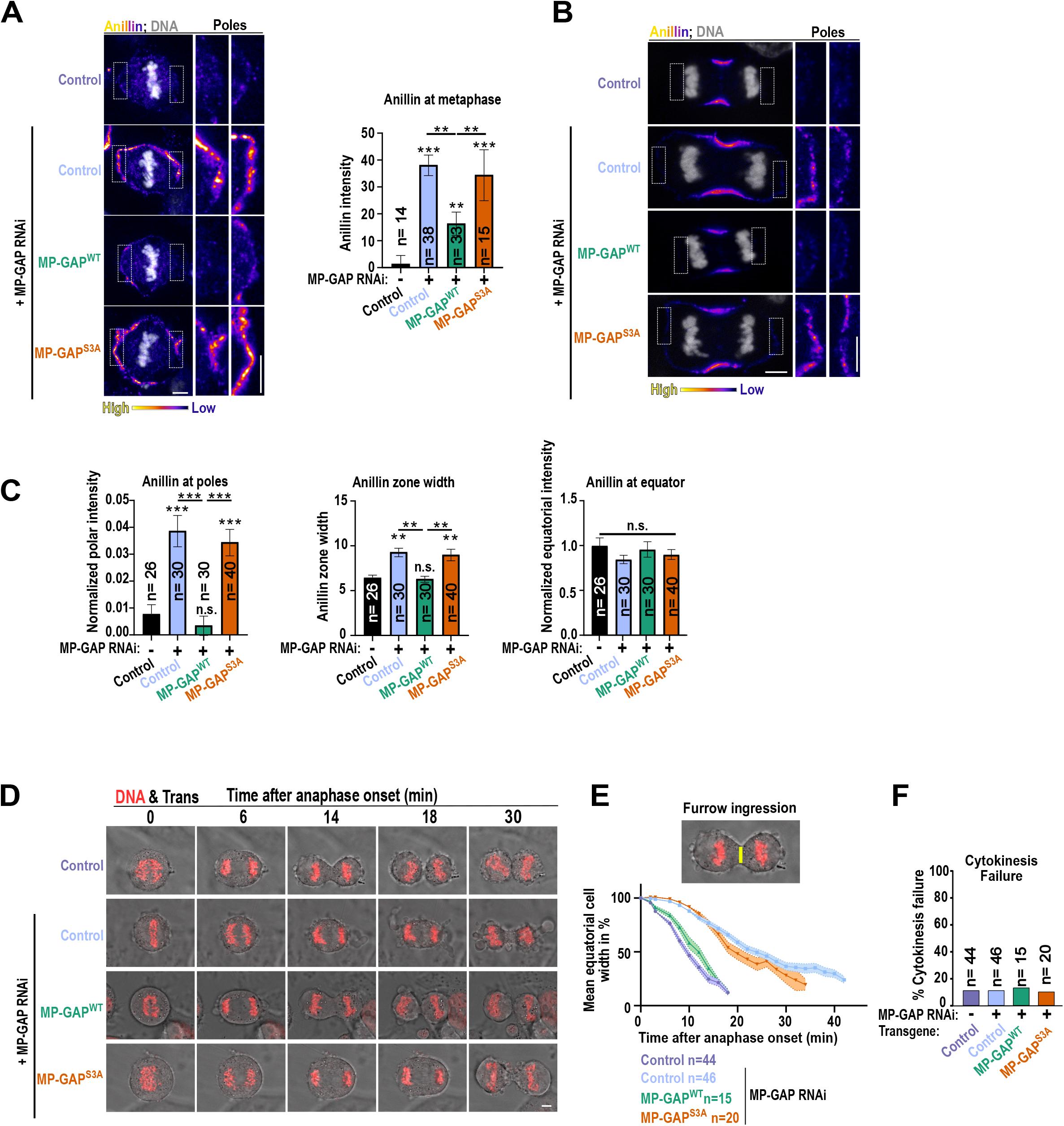
Aurora A kinase phosphorylation of MP-GAP limits anillin accumulation to a narrow equatorial zone and at the poles of the cell. A) Shown are immunofluorescence images of anillin (pseudocolour) and DNA (grey) stained metaphase HeLa cells. Control cells (no transgene), GFP-MP-GAP^WT^ and GFP-MP-GAP^S3A^ expressing cells were treated with or without MP-GAP siRNA as indicated. Magnifications of the polar cortex are shown (left). Mean cortical anillin fluorescence intensity for indicated conditions (right). B) Shown are immunofluorescence images of anillin (pseudocolour) and DNA (grey) stained anaphase HeLa cells for the indicated conditions. Magnifications of the polar cortex are shown. Intensity scalings are increased for the magnifications to better visualize polar anillin accumulation. C) Plotted is the mean anillin fluorescence intensity the cell poles, at the cell equator and the zone width in anaphase for the indicated cell lines and RNAi conditions. D) Merged time-lapse transmission and DNA (SiR-DNA, red) images of HeLa cells expressing no transgene (control), MP-GAP^WT^ or MP-GAP^S3A^ for the indicated RNAi conditions. E) Plotted is the mean equatorial cell width over time for the indicated conditions. F) Plotted is the % of cytokinesis failure during live-cell imaging for the indicated conditions. For all n = number of cells, and ≥2 independent experiments were performed for each condition. Error bars are SEM, and p values were calculated with the Kruskal-Wallis test. Scale bars are 5 µm.

Lastly, we tested whether MP-GAP depletion has any effect on furrow ingression. We filmed HeLa cells after MP-GAP depletion and measured the equatorial and pole-to-pole cell width from anaphase onset until the contractile ring was fully ingressed. In control cells, the equatorial cell width decreased and the pole-to-pole cell width increased after anaphase onset (Fig. 3D, E, S2C). Both, the decrease in equatorial cell width and the increase in pole-to-pole cell width was strongly delayed in cells depleted of MP-GAP by RNAi, but eventually most cells completed cytokinesis (Fig. 3D-F, S2C). Expression of MP-GAP^WT^, but not MP-GAP^S3A^ rescued the delay in furrow formation and ingression and pole-to-pole cell elongation.

Altogether, after MP-GAP depletion cortical anillin levels at the cell poles increased and the equatorial anillin zone widened, which resulted in delayed cell elongation and furrow ingression. Expression of wild-type MP-GAP, but not the Aurora A kinase phosphodeficient mutant, rescues the MP-GAP depletion phenotype suggesting that phosphorylation of MP-GAP by Aurora A kinase promotes MP-GAP activity.

### F-actin, but not Aurora A phosphorylation, targets MP-GAP to the cell periphery

Aurora A kinase localizes to the centrosomes and astral microtubules (Kufer et al., 2002; Portier et al., 2007; Stenoien et al., 2003; Sugimoto et al., 2002; Ye et al., 2015) and MP-GAP to the cell periphery (Zanin et al., 2013). Therefore, phosphorylation of MP-GAP by Aurora A could occur around the spindle poles and/or at the cell periphery. To analyze whether and where MP-GAP and Aurora A are in close proximity within the cell we performed in situ proximity ligation assays (PLA). Cells were incubated either with anti-Aurora A and anti-MP-GAP primary antibodies or only with one primary antibody as a negative control. If both epitopes are less than 10 nm apart bright PLA foci are expected. Indeed, we observed numerous PLA foci when cells were incubated with both primary antibodies but not in the negative control (Fig. 4A). This suggests that Aurora A and MP-GAP are in close proximity and might interact with each other. To analyze the subcellular distribution of the PLA foci within the cell, we split the cell into three regions: periphery, middle and center and counted the number of foci in each region. PLA foci were strongly enriched at the cell periphery in comparison to the middle or center in metaphase and anaphase (Fig. 4B). Lastly, we determined whether PLA foci were equally present at the cell equator and poles in anaphase cells. We found more foci enriched at the polar in comparison to the central area (Fig. 4C).

**Figure 4.**
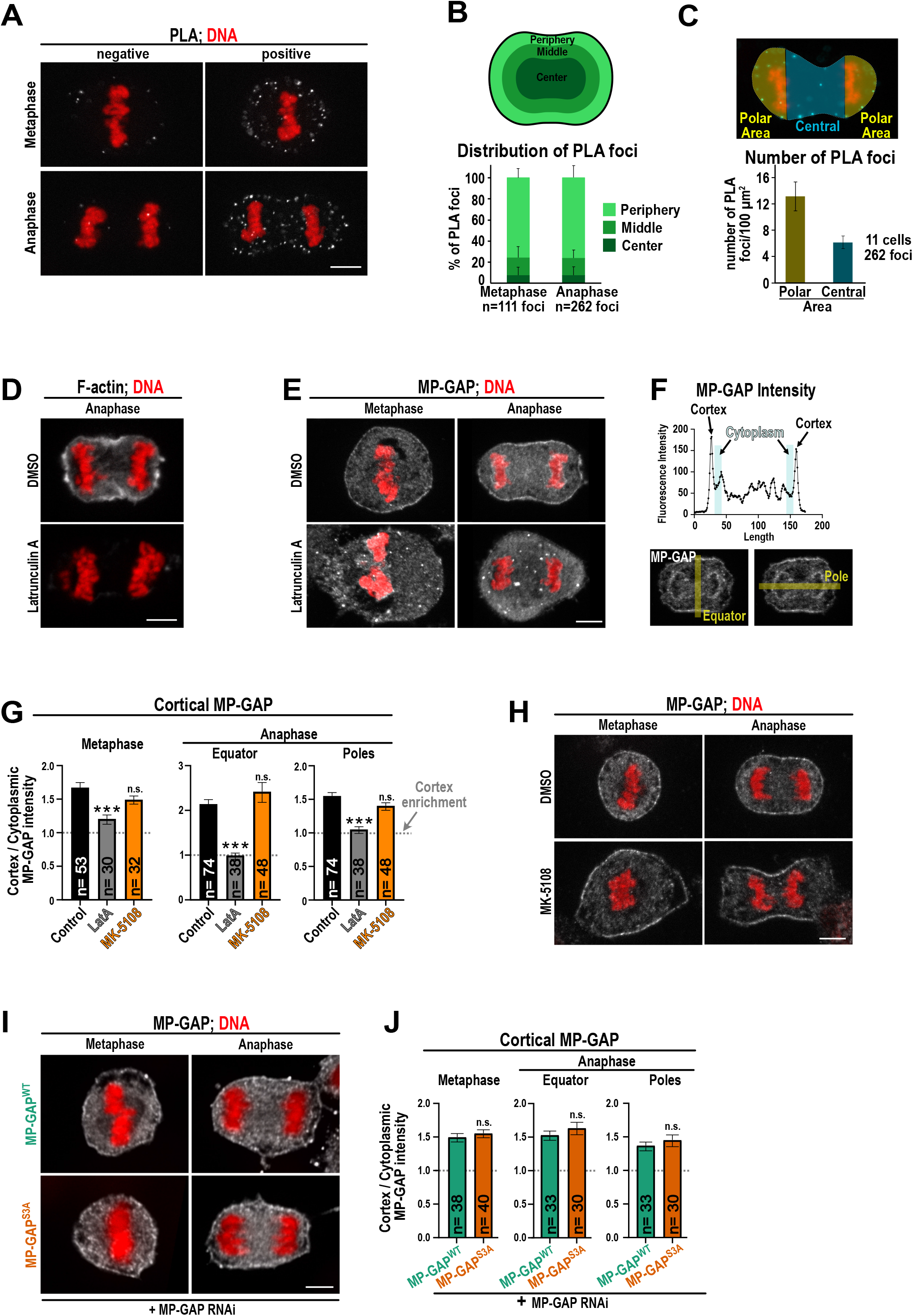
MP-GAP is targeted to the actin cortex independently of Aurora A activity. A) Merged immunofluorescent images showing PLA foci (white) and DNA (red) for metaphase and anaphase HeLa cell. Staining was performed either with the anti-Aurora A and anti-MP-GAP together (positive) or only with one antibody (negative). B) Distribution of the PLA foci in % at the cell periphery, middle or center during metaphase and anaphase. C) The number of PLA foci per 100 µm^2^ is plotted for the central or polar area of the cell. D) Immunofluorescence images of F-actin (white) and DNA (red) stained HeLa cells treated with DMSO or 2.5 µM Latrunculin A (LatA). E) Immunofluorescence images of MP-GAP (white) and DNA (red) stained HeLa cells treated with DMSO or 2.5 µM Latrunculin A. F) MP-GAP fluorescence intensity is plotted for an equatorial linescan. Examples of an equatorial and polar linescan are shown for illustration. G) Quantification of the mean cortical MP-GAP intensity as depicted in (F) for the entire cell cortex in metaphase and for the poles and equator in anaphase cells for indicated treatments. Values above the dotted line represent cortical MP-GAP enrichment. H) Shown are immunofluorescence images of metaphase and anaphase HeLa cells treated with DMSO or MK-5108. Cells were stained for MP-GAP (white) and DNA (red). I) Shown are immunofluorescence images of metaphase and anaphase HeLa cells expressing GFP-MP-GAP^WT^ or GFP-MP-GAP^S3A^ and treated with MP-GAP siRNA. Cells were stained for MP-GAP (white) and DNA (red). J) Quantification of the mean cortical MP-GAP intensity as depicted in (F) for the entire cell cortex in metaphase and for the poles and equator in anaphase cells for indicated treatments. For all n = number of cells, and ≥2 independent experiments were performed for each condition. Error bars are SEM, and p values were calculated with the Kruskal-Wallis test. Scale bars are 5 µm.

MP-GAP localizes to the cell periphery in metaphase and anaphase (Zanin et al., 2013) and in *C. elegans* the MP-GAP homologue requires F-actin polymerization for cortical targeting (Michaux et al., 2018). Thus, we asked whether cortical localization of human MP-GAP also depends on the presence of F-actin. Inhibition of F-actin polymerization by Latrunculin A treatment caused a strong reduction of cortical F-actin and MP-GAP during metaphase and anaphase (Fig. 4D, E). To quantitatively measure changes in MP-GAP localization, we drew linescans across the polar and equatorial regions of the cells and determined the enrichment of MP-GAP at the cell periphery (Fig. 4F). In control metaphase cells MP-GAP is ∼1.5 fold enriched at the cortex in comparison to the cytoplasm. During anaphase MP-GAP exhibits ∼2 fold enrichment at the cell equator and a ∼1.5 enrichment at the cell poles. After F-actin depolymerization by Latrunculin A addition, MP-GAP enrichment at the entire cortex is lost and equals cytoplasmic levels (Fig. 4G).

Since Aurora A preferentially phosphorylates MP-GAP at the cell periphery we asked whether Aurora A phosphorylation of MP-GAP is required for MP-GAP targeting to the cell periphery. After Aurora A inhibition cortical MP-GAP enrichment was not reduced in comparison to DMSO treated control cells (Fig. 4G, H). This observation was supported by quantitatively measuring cortical localization of MP-GAP^WT^ and MP-GAP^S3A^ after the depletion of endogenous MP-GAP. MP-GAP^WT^ and MP-GAP^S3A^ enriched to a similar extent at the cell cortex in metaphase and anaphase (Fig. 4I, J).

Concluding, MP-GAP localization to the cell cortex requires F-actin polymerization. Although, Aurora A phosphorylates MP-GAP preferentially at the cell periphery in mitotic cells and particularly at the polar area in anaphase cells, Aurora A is not involved in targeting MP-GAP to the actin cortex.

### The F1 region of MP-GAP binds the catalytic GAP domain

MP-GAP consists of the N-terminal localized catalytic GAP domain and a large C-terminal region of unknown function. To check whether MP-GAP could adopt an autoinhibitory conformation, we tested binding of the F1 or F2 region of MP-GAP to the catalytic GAP domain. We immobilized purified GST-tagged GAP, F1 or F2 MP-GAP fragments on beads and incubated them with His-tagged GAP domain. We observed that GST-tagged F1, but not F2 or the GAP domain itself, bound to the His-tagged GAP domain (Fig. 5A). This points toward the possibility that the binding of F1 to the GAP domain inhibits GAP activity. Since Aurora A phosphorylates the F1 region, this could regulate binding of F1 to the GAP domain. If the phosphorylation sites are in the binding surface, interaction between F1 and the GAP is expected to be abolished. If the phosphorylation sites are in the hinge region connecting the catalytic GAP domain with the binding surface in F1, binding should still occur. To test this, we generated a phosphomimetic F1 (F1^S3D^) region and tested for GAP binding. The phosphomimetic F1^S3D^ fragment still bound the GAP domain (Fig. 5B), indicating that the Aurora A target sites are in the hinge region that connects the GAP domain with the F1 fragment. Together, the F1 region binds the GAP domain suggesting that MP-GAP adopts an autoinhibitory conformation.

**Figure 5.**
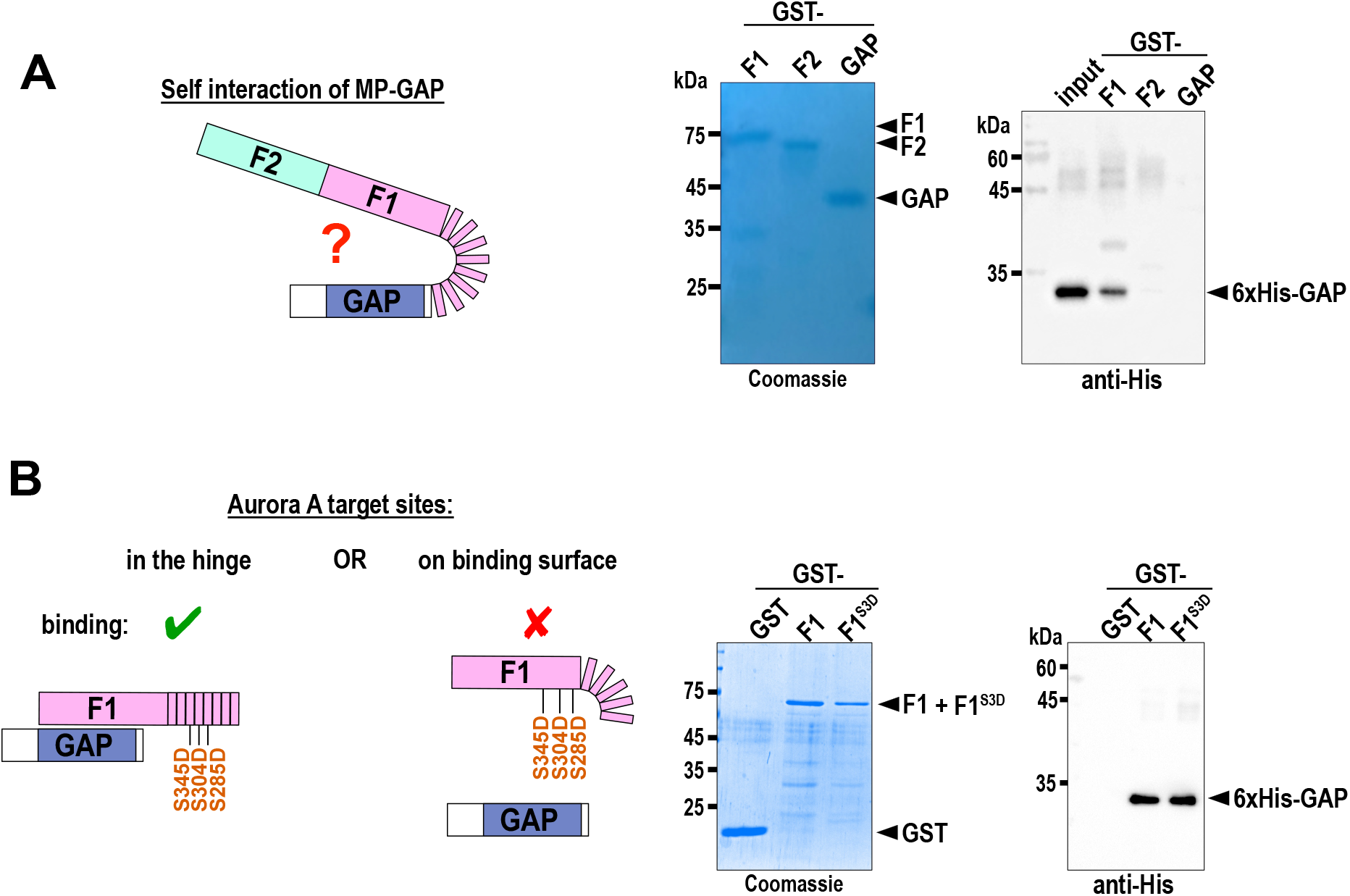
The F1 region binds the catalytic GAP domain of MP-GAP. A) Scheme of the possible autoinhibitory conformation of MP-GAP (left). Coomassie gel and anti-His probed immunoblot of pull down assays incubating immobilized GST-tagged MP-GAP fragments (F1, F2 and GAP) with soluble 6xHis-tagged GAP domain (right). B) Scheme of the putative position of the Aurora A phosphorylation sites in either the hinge region (left) or on the binding surface (right) of F1. Coomassie gel and anti-His probed immunoblot of pull down assays incubating immobilized GST-tagged F1 wild type or F1^S3D^ with soluble 6xHis-tagged GAP domain (right).

### Aurora A-dependent signaling and Ect2 autoinhibition protect the cell against cytokinesis failure

Aurora A inhibition did not cause cytokinesis failure although more anillin accumulated at the cell poles and the equatorial anillin zone was broader. We asked whether Aurora A inhibited cells succeed in cytokinesis since RhoA activation hardly occurs outside the cell equator due to the autoinhibition of Ect2 (Fig. 6A). Ect2 autoinhibition by the BRCT2 domain is only released at the cell equator which results in local RhoA activation (Chen et al., 2019; Schneid et al., 2021) (Fig. 6A). Mutating W307 of the BRCT2 domain to alanine abolished Ect2 autoinhibition and increased RhoA signaling but did not cause cytokinesis failure (Chen et al., 2019; Schneid et al., 2021), most likely, since Aurora A activity still promotes RhoA inactivation at the cell poles. To test whether simultaneous inhibition of Aurora A and absence of Ect2 autoinhibition cause cytokinesis failure, we employed a previously established genetic replacement system for Ect2, in which RNAi resistant Ect2^WT^ and Ect2^W307A^ are expressed in HeLa cells (Schneid et al., 2021). We inhibited Aurora A kinase during live-cell imaging before anaphase onset in Ect2^WT^ and Ect2 ^W307A^-expressing cells after the depletion of endogenous Ect2 (Fig. 6B). Similar to HCT-116 cells, Aurora A inhibition in Ect2^WT^ HeLa control cells did not elevate cytokinesis failure rate (Fig. 6B-D). In contrast, Aurora A inhibition in Ect2^W307A^-expressing cells strongly increased cytokinesis failure rate and the percentage of cells with shape oscillation (Fig. 6B-D, S3A, B).

**Figure 6.**
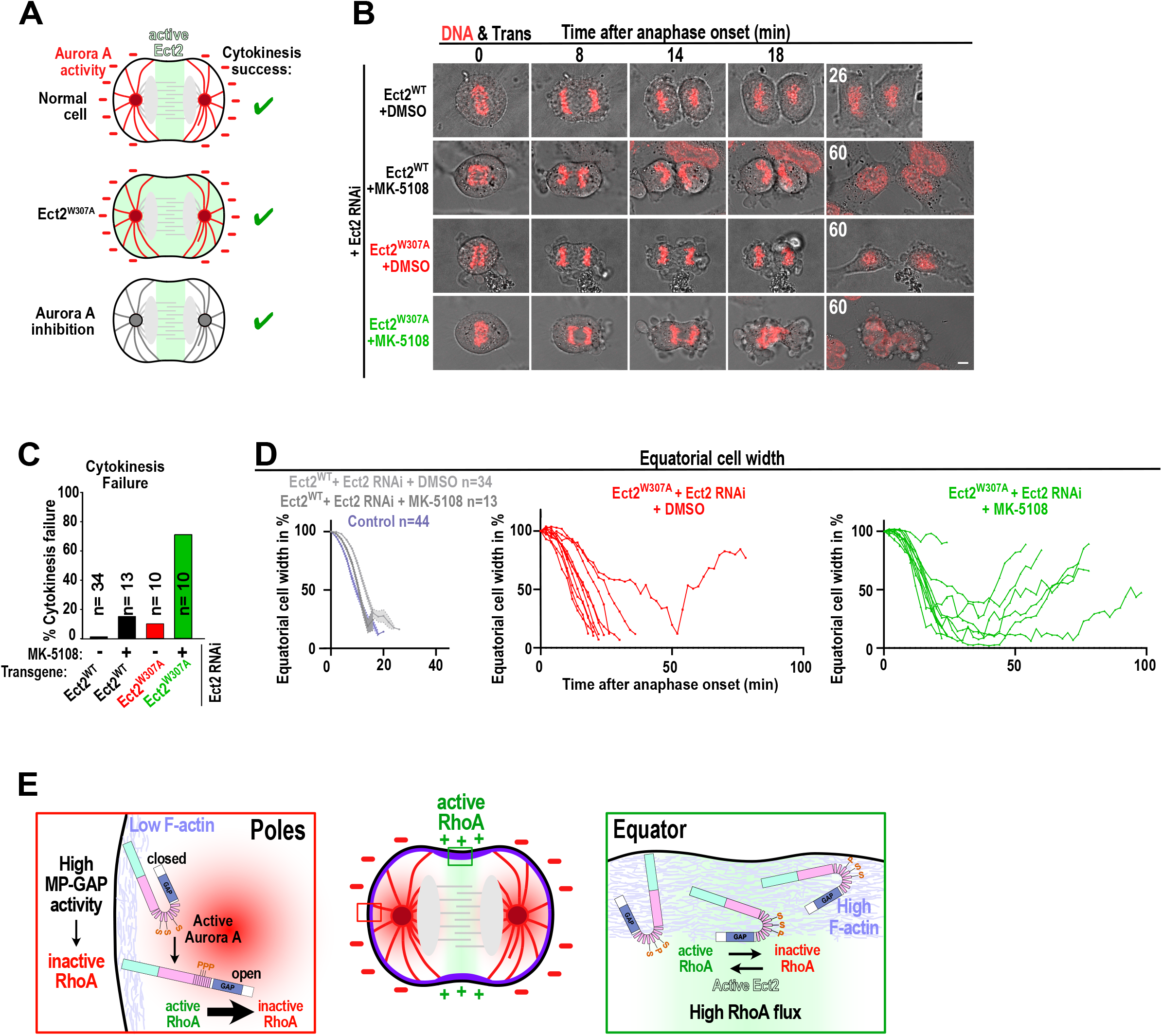
Autoinhibition of Ect2 and Aurora A kinase-signaling protect the cell against cytokinesis failure. A) In normal cells Aurora A signal is high at the poles and Ect2 autoinhibition is locally released at the cell equator (top). Abolishing Ect2 autoinhibition by the W307A mutation, results in active Ect2 throughout the cell (middle). Without Aurora A signaling, Ect2 autoinhibition is still locally released at the equator (bottom). In all conditions cytokinesis succeeds. B) Merged time-lapse transmission and DNA (SiR-DNA, red) images of HeLa cells expressing Ect2^WT^ or Ect2^W307A^. Cells were treated during live-cell imaging around anaphase onset with DMSO or MK-5108 as indicated. C, D) Percentage of cytokinesis failure (C) and equatorial cell width (D) of the conditions filmed in (B). E) Two spatially distinct regulatory inputs control MP-GAP function at the cell equator and the cell poles during anaphase. At the cell poles F-actin levels are low, but Aurora A kinase activity is high. Aurora A phosphorylates the hinge region between the GAP domain and the F1 region resulting in opening and activation of MP-GAP and RhoA inactivation. At the cell equator Ect2 autoinhibition is released, RhoA is activated and promotes F-actin polymerization. Due to high F-actin levels MP-GAP enriches at the equator and increases the GTP hydrolysis rate of RhoA resulting in high RhoA flux and narrowing of the active RhoA zone. For all n = number of cells, and ≥2 independent experiments were performed for each condition. Scale bars are 5 µm.

In summary, polar inhibition of RhoA activity mediated by Aurora A signaling and Ect2 autoinhibition are redundant mechanisms to spatially pattern RhoA activity and protect against cytokinesis failure.

## Discussion

During cytokinesis stimulatory and inhibitory signals spatially control RhoA activity and thereby induce contractile ring formation and cleavage furrow formation at the right location. Here we reveal that the activity and localization of the negative regulator of RhoA, MP-GAP, is controlled by two distinct regulatory inputs which together spatially limit RhoA signaling during cytokinesis. Based on our data we suggest the following model: MP-GAP activity could be suppressed by the binding of the F1 region to the catalytic GAP domain. This autoinhibition is released by the phosphorylation of three amino acid residues by Aurora A kinase in the hinge region between the F1 region and the GAP domain. Since Aurora A is active in a gradient around spindle-poles this results in a preferential increase of MP-GAP activity at the overlaying polar cortex and consequently RhoA inactivation at the polar membrane (Fig. 6E). Our findings identify a long-searched spindle pole-located inhibitory signal suppressing RhoA activity at the cell poles. However, MP-GAP restricts RhoA activity not only at the cell poles but also at the cell equator. MP-GAP requires an intact F-actin cortex to enrich at the cell periphery and since F-actin levels are high at the equator, total cortical MP-GAP levels are also higher at the equator compared to the poles of the cell. At the equator RhoA is continuously activated by Ect2 and together with MP-GAP-mediated RhoA inactivation this results in a high RhoA flux through the GTPase cycle and thereby restricts active RhoA to a narrow equatorial zone. Collectively, MP-GAP activity integrates two regulatory inputs: targeting to the cell periphery by F-actin and elevation of MP-GAP activity by Aurora A phosphorylation (Fig. 6E). Together both inputs allow independent fine tuning of the amounts and activity of MP-GAP: if RhoA activity is high, F-actin polymerization is induced and MP-GAP is recruited and increases RhoA flux to form a narrow active RhoA zone. At the cell poles where F-actin levels and subsequently MP-GAP levels are low, MP-GAP activity is boosted by Aurora A phosphorylation to shut RhoA off.

Aurora A turns over rapidly (Portier et al., 2007; Stenoien et al., 2003) and is active in a gradient around monopolar spindles (Ye et al., 2015). Our data show, that Aurora A preferentially interacts with MP-GAP at the polar cell periphery and not at the spindle poles, suggesting that active Aurora A diffuses from the spindle poles to the overlying cortex where it phosphorylates F-actin-bound MP-GAP. Using pull down assays we demonstrate that the F1 region of MP-GAP binds the catalytic GAP domain which points toward the possibility that MP-GAP adopts an autoinhibitory conformation. Intriguingly, the three identified Aurora A target sites are located in the F1 fragment suggesting that phosphorylation of those sites regulates F1 binding to the GAP domain. Exchanging Aurora A phosphorylation sites to phosphomimetic residues did not block binding of F1 to the GAP domain and therefore, we speculate that the Aurora A target sites are located in the hinge between the binding surface of the F1 region and the GAP domain. Phosphorylation of the hinge region by Aurora A might promote opening and increase GAP activity.

Aurora A inhibition or MP-GAP depletion increases cortical anillin levels not only during anaphase but also in metaphase. Thus, Aurora A elevates MP-GAP activity already in metaphase. In metaphase the cell is round and the spindle poles are closely together and therefore a uniform Aurora A activity throughout the cell is expected. The metaphase signaling of Aurora A ensures, together with CDK1 phosphorylation of Ect2 (Su et al., 2011), that RhoA activity remains low prior to anaphase onset. Elevated RhoA activity levels during metaphase cause spindle positioning and chromosome segregation defects and might be responsible for the protumorigenic activity of RhoA (Limzerwala et al., 2020).

During anaphase the cell elongates and the distance between the two spindle poles increases. Due to its centrosome-centered localization, Aurora A activity is expected to be graded, high at the cell poles and low at the cell equator. Thereby MP-GAP activation at the cortex is particularly enforced at the cell poles. However, MP-GAP is not only present at the polar cortex but also enriches at the equatorial cortex. MP-GAP requires F-actin to localize to the cell cortex demonstrating that MP-GAP F-actin binding or F-actin associated protein binding is conserved from worms to humans (Michaud et al., 2022; Michaux et al., 2018). Since active RhoA and consequently F-actin levels are high at the cell equator, MP-GAP enriches there and has some basal GAP activity. Active RhoA is tethered to the plasma membrane with its lipid moiety and is highly dynamic (Budnar et al., 2019). Therefore, active RhoA is predicted to diffuse away from the site of activation and distribute over the plasma membrane. However, this is not observed in cells and therefore, it was suggested that inactivation of RhoA and accelerating RhoA flux through the GTPase cycle by the RhoA GAP is prerequisite for the formation of a narrow active RhoA zone at the cell equator (Bement et al., 2006). Indeed MP-GAP depletion causes a broader anillin zone, which suggests that equatorial MP-GAP-accelerated RhoA flux restricts RhoA zone dimensions. In the presented study, the efficiency of MP-GAP depletion was improved in comparison to our previous publication (Zanin et al., 2013), which explains the stronger phenotypes. A phosphodeficient MP-GAP mutant or Aurora A inhibition also broadened the anillin zone suggesting that Aurora A phosphorylation, to some extent, also activates MP-GAP at the cell equator. Consistently, we also observed some MP-GAP-Aurora A PLA foci at the equatorial cortex, although their number was reduced in comparison to the cell poles. Spatially altering RhoA activity during anaphase by MP-GAP depletion or abolishing Aurora A phosphorylation of MP-GAP strongly delayed pole-to-pole elongation and furrow ingression demonstrating that spatially limiting RhoA activity is needed for the overall cell shape changes during cytokinesis.

Abolishing Aurora A signaling and Ect2 autoinhibition together resulted in high frequency of cytokinesis failure suggesting that limiting RhoA activity by one mechanism is sufficient however if both are absent cytokinesis fails. After the disruption of Aurora A signaling (this work) or Ect2 autoinhibition (Schneid et al., 2021) a wider anillin zone still forms. This is possible since both pathways use spatial landmarks of the mitotic spindle to pattern RhoA activity: Aurora A signaling is high above the spindle poles where it shuts off RhoA activity and Ect2 adopts an autoinhibitory conformation, which is only released at the spindle midzone. In the absence of Aurora A signaling, Ect2 autoinhibition is still preferentially released above the midzone and therefore the RhoA zone still forms at the cell equator although it is broader. In case Ect2 autoinhibition is lost, Aurora A provides a spatial clue for RhoA inactivation at the cell poles. Together these two complementary mechanisms of controlling RhoA activity are the foundation for the enormous robustness of cytokinesis.

Recent studies revealed two mechanistically distinct pathways that limit the accumulation of myosin II and F-actin on the poles in an astral microtubule-dependent manner. In *C. elegans* polar myosin II is removed by dynein (Chapa-y-Lazo et al., 2020) and in human tissue culture cells polar y-actin nucleation is blocked by the inhibition of formin (Chen et al., 2020). Both mechanisms do not block RhoA activity directly and thus might become essential when MP-GAP-mediated RhoA inactivation at the cell poles is compromised. In addition, several spindle-pole independent signals emanating from the spindle-midzone, the kinetochores and the chromatin (Beaudet et al., 2017; Beaudet et al., 2020; Kiyomitsu and Cheeseman, 2013; Ramkumar et al., 2021; Rodrigues et al., 2015) control the dynamics of contractile ring components at the cell poles. For example, spindle midzone enriched Aurora B promotes F-actin removal from the cell poles (Ramkumar et al., 2021). We did not observe an increase of anillin at the poles or a wider zone after Aurora B inhibition, similar to previous observations (Adriaans et al., 2019). This suggests that Aurora B has a more direct role in blocking F-actin nucleation at the polar cortex maybe by regulating F-actin nucleators or stabilizers. Future investigations will need to study whether those pathways work redundantly with Aurora A signaling and whether specific cell types rely on selected pathways. A recent study suggests that AIR-1, the Aurora A homologue in *C. elegans*, prevents ECT-2 membrane binding by phosphorylating the PH domain (Longhini and Glotzer, 2022). Whether Aurora A controls Ect2 localization and/or activity in addition to MP-GAP in human cells must be studied in the future. In summary, our work provides mechanistic insights into how the regulation of MP-GAP activity and localization by Aurora A and F-actin patterns RhoA activity during mitosis. Spatially and temporally restricted small GTPases activity zones are frequently observed during many cellular processes including cell migration, cell polarization or wound healing (Bement and von Dassow, 2014). Thus, the modulation of GAP localization and activity by distinct spatial clues might also shape GTPase activity zones in other cellular contexts. Furthermore, Aurora A kinase also inhibits contractility during other cellular processes such as cell polarization (Kapoor and Kotak, 2019; Klinkert et al., 2019; Zhao et al., 2019) where the same pathway might act as well. Aurora A kinase is also an oncogene and overexpression has been reported in a broad range of human malignancies (Du et al., 2021), thus, a better understanding of Aurora A’s role in cell division might be crucial for the development of new therapeutic treatments.

## Material and Methods

### Cell culture

Human HCT-116 and HeLa Flp-In T-Rex cells were grown at 37°C in 5% CO_2_ in McCoy 5A growth media (for HCT-116, Thermo Scientific, 16600082) and Dulbecco’s growth medium (for HeLa, DMEM, Thermo Scientific, 61965059) supplemented with 10% FBS (Thermo Scientific, 10270106) and 1% penicillin/streptomycin (PS, Merck, A2213). CRISPR-tagged Neongreen-Anillin HCT-116 cell line was a gift from Alisa Piekny (Husser et al., 2022). Isogenic HeLa cell line harboring stable integration of RNAi resistant GFP-tagged wild-type MP-GAP was used from a previous study (Zanin et al., 2013). GFP-tagged MP-GAP^S3A^ (S285A, S304A, and S345A) phosphodeficient transgene was obtained by FRT/Flp-mediated recombination. HeLa Flp-In host cell was transfected with pOG44 and GFP-MP-GAP^S3A^ using X-tremeGene 9 (Sigma Aldrich, 8079398949). After transfection, cells were cultured in DMEM (10% FBS, 1% PS) supplemented with hygromycin (0.2 mg/ml) to select for transgene integration. After 2.5 weeks the clones were pooled. In all MP-GAP^WT^ and MP-GAP^S3A^ experiments, transgene expression was induced with 0.2 µg/ml tetracycline.

### Cloning of MP-GAP variants

The MP-GAP fragments used for pulldown (GST-GAP: AA2-253 (pEZ365); GST-F1 AA255-641 (pEZ359); GST-F2 AA641-1023 (pEZ360)), or antibody generation (GST-MP-GAP^AA560-801^ (pEZ229)) were cloned into the pGEX-4T using Gibson assembly. The phosphodeficient mutants (GST-F1-C1: S285A, S304A, and S345A (pEZ371)) and single site mutations (S304A (pEZ366), S345A (pEZ367), S345A (pEZ368)) of F1 were obtained by side-directed mutagenesis. The GST-F1-C2 mutant (S403A, S434A, S500A and T508A (pEZ377)) was synthesized by Eurofins and subcloned into the pGEX-4T vector with Gibson assembly. The phosphomimetic F1 fragment (GST-F1-S3D, S285D, S304D, and S345D (pEZ426)) was generated by site-directed mutagenesis and cloned into pGEX-4T. RNAi resistant MP-GAP^S3A^ (S285A, S304A, and S345A, pEZ378) was cloned into the pcDNA5/FRT/TO vector containing a GFP-tag using Gibson assembly. All plasmids generated for the study are listed in Table 1.

**Table 1.** Plasmids generated for the study.

| **Plasmid number** | **Content** |
| --- | --- |
| pEZ229 | GST-MP-GAP AA560-801 |
| pEZ359 | GST-F1 MP-GAP AA255-641 |
| pEZ360 | GST-F2 MP-GAP AA641-1023 |
| pEZ365 | GST-GAP MP-GAP AA2-253 |
| pEZ366 | GST-F1 MP-GAP AA255-641 (S285A) |
| pEZ367 | GST-F1 MP-GAP AA255-641 (S304A) |
| pEZ368 | GST-F1 MP-GAP AA255-641 (S345A) |
| pEZ371 | GST-F1-C1 MP-GAP AA255-641 (S285A, S304A, S345A) |
| pEZ377 | GST-F1-C2 MP-GAP AA255-641 (S403A, S434A, S500A and T508A) |
| pEZ378 | DNA5/FRT/TO GFP-MP-GAP<sup>S3A</sup> (S285A, S304A, S345A) |
| pEZ426 | GST-F1 MP-GAP AA255-641 (S285D, S304D, S345D) |

### siRNA transfection

The phenotype of MP-GAP^WT^ and MP-GAP^S3A^ was determined after depleting endogenous MP-GAP by RNAi. One day prior to siRNA transfection cells were plated in 12 well plates with DMEM (10%FBS, 0.2 mg/ml tetracycline) at approximately 30% confluency. For immunofluorescence, cells were seeded on 18 mm poly-L-lysine (P1524-100G, Sigma) coated glass coverslips (P233.1, Menzel) and for live-cell imaging cells were seeded into ibidiTreat chambers (80426, Ibidi). Next day, cells were transfected with a final concentration of 40 nM MP-GAP siRNA oligo (ATTACAGGCTGCAGTAGTA, J-021122-05-0020, Dharmacon, (Zanin et al., 2013)) using Interferin (Polyplus-transfection, 101000028). 18-20 hr after siRNA transfection 0.5 mM thymidine (T1895, Sigma) was added to synchronize the cell cycle. After 22-24 hr incubation, cells were released from the thymidine block by washing 3x for 5 min with DMEM (10% FBS, 0.2 mg/ml tetracycline). 8-9 hr after the thymidine release live-cell imaging was started or cells were fixed for immunofluorescence of mitotic cells.

Same procedure was followed for depletion of endogenous Ect2 (Fig. 6), however, cells were transfected with a final concentration of 30 nM Ect2 siRNA oligo (GGCGGAAUGAA CAGGAUUU, Dharmacon, (Yüce et al., 2005)) using Lipofectamine RNAiMAX (13778075, Thermo Fischer).

### MP-GAP antibody production

Polyclonal antibodies against purified GST-tagged MP-GAP^AA^ ^560-801^ were generated in rabbits. GST-MP-GAP^AA560-801^ expression was induced in BL21(DE3) *E. coli* for 4 hr at 37°C with 0.5 mM IPTG and purified over a glutathione-Sepharose column. After depleting GST antibodies from the obtained rabbit serum, the MP-GAP antibodies were purified with GST-MP-GAP^AA^ ^560-801^-coupled SulfoLink resin. Antibodies bound to the SulfoLink resin were eluted in 0.2 M glycine pH 2.5 and neutralized by 1 M TrisHCl pH 8.5. Antibodies were concentrated and the buffer was exchanged to 1x Phosphate Buffered Saline (PBS) using Pierce Protein Concentrator (10K MWCO; Life Technologies, 88527). Specificity of the MP-GAP antibody was verified by the absence of immunostaining after MP-GAP depletion by RNAi.

### Immunostaining

For anillin stainings cells were fixed in ice-cold 10% TCA (Merck, 1008070100) or 100% ice-cold methanol for 15 min on ice. For MP-GAP stainings cells were fixed in 100% ice-cold methanol. After fixation cells were washed in 1x PBS and incubated with blocking solution (1x PBS, 4% BSA, 0.1% Triton) for 1 hr at room temperature (RT). As primary antibodies anti-anillin (1:3000, self-made), or MP-GAP (1:250, self-made) were diluted in blocking solution and incubated for 2 hr at RT. Coverslips were washed 3x 5 min in 1x PBS/0.1% Triton and incubated with Alexa FluorTM 568 donkey anti-rabbit (1:500, A10042, Life Technologies) and Alexa FluorTM 635 goat anti-mouse (1:500, A31574, Life Technologies), and Hoechst 33258 (861405; Sigma-Aldrich) in blocking solution for 1 hr at RT.

For visualizing astral microtubules after MK-5108 treatment in HCT-116 cells, α-tubulin (1:10 000, T6199, Sigma) stainings were performed by washing coverslips once with 1x PBS and fixing in 1x PBS containing 0.5% glutaraldehyde (340855, Sigma), 0.5% Triton for 10 min at RT. After fixation coverslips were washed 3x 10 min with freshly made 1x PBS containing 0.1% NaBH4 (806373, Merck) at RT. Cells were washed once with 1x PBS, blocked and stained for α-tubulin and DNA as described above. For F-actin staining after Latrunculin A treatment, cells were fixed in 4% cold PFA. Following fixation, slides were washed in 1x PBS and incubated with blocking solution (1x PBS, 4% BSA, 0.1% Triton) for 1 hr at RT. After blocking, slides were incubated with Alexa Fluor™ 568 Phalloidin (1:500, Life Technologies, A12380) and Hoechst 33258 (861405; Sigma) in blocking solution for 1 hr at RT. For all stainings: at the end coverslips were washed 3x 5 min in 1x PBS/0.1% Triton and covered with mounting medium (1x PBS, 4% n-Propyl-Gallate, 90% Glycerol) and stored at −20°C.

### Immunoblotting

For immunoblotting cell lines were seeded in 6-well plates, transfected with MP-GAP siRNA, and synchronized with single thymidine block as described above. Cells were collected 8-9 hr after thymidine release. Samples were heated at 95°C for 5 min and sonicated in a water bath for 20 min. As primary antibodies rabbit anti a-actin (1:3000, A-1978, Sigma) and rabbit anti-MP-GAP (1:100, self-made) were used; and as secondary antibody HRP-conjugated anti-rabbit or anti-mouse (both 1:10 000; anti-rabbit (1706515; Bio-Rad); anti-mouse (170-6516; Bio-Rad) were used. Membranes were developed with ECL^TM^ Prime Western Blotting (Sigma, Cytiva RPN2236). Expression levels of the endogenous and the GFP-tagged MP-GAP proteins were quantified by measuring the intensities of the MP-GAP and a-actin bands using the ImageJ software. GFP-tagged and endogenous MP-GAP levels were first normalized to a-actin levels and subsequently the levels of GFP-tagged MP-GAP variants relative to endogenous MP-GAP were calculated.

### Inhibitor treatments with MK-5108, AZD1152-HQPA and Latrunculin A

Stock solutions MK-5108 (Selleckchem, S2770, 10 mM), AZD1152-HQPA (Sigma, SML0268-10MG, 10 mM) and Latrunculin A (Enzo, BML-T119-0100, 10 mM) were made in DMSO and stored at −20°C. Inhibitors were diluted in complete cell culture medium and used at the following final concentrations: 10 µM for MK-5108, 200 nM for AZD and 2.5 µM for Latrunculin A. For immunostaining cells were synchronized with a single thymidine block and 8 hr after release they were treated for 10-24 min with 10 µM MK-5108 and 200 nM AZD, or for 20 min with 2.5 µM Latrunculin A. For live-cell imaging, MK-5108 and AZD were present on the cells for maximum 26 min and 14 min, respectively.

### Aurora A kinase assays

Aurora A kinase assays were performed in kinase buffer (25 mM HEPES pH 7.5, 50 mM NaCl, 10 mM MgCl2, 0.1% Triton X-100 and 1 mM DTT (Gomez-Cavazos et al., 2020) containing 300 ng GST-tagged MP-GAP fragments and 200 ng recombinant human Aurora A kinase (Merck, 14-511), 1mCi of [gamma-P32]ATP (Hartmann Analytic, SRP-801) and 0.2 mM dATP (mi-N1010L, Metabion) with a final volume of 20 µl. The kinase reaction was incubated at 30°C for 30 min and it was terminated by the addition of 2x sample buffer (120 mM Tris pH 6.8, 4% SDS, 20% Glycerol, 0.02% bromphenol blue, 5% b-mercaptoethanol). Samples were heated to 95°C for 5 min and then separated by SDS-PAGE. The SDS-Gel was stained with Coomassie blue (B0149, Sigma) and dried prior to autoradiography on Whatman filter paper. Phosphor imaging was performed with a Typhoon Variable Mode Imager (Amersham Biosciences) connected to Amersham^TM^ Typhoon^TM^ scanner control software.

### Proximity ligation assay

Proximity ligation assays with Aurora A kinase and MP-GAP were performed in HeLa cells using mouse anti-Aurora A Kinase antibody (1:500, A1231 Merck), rabbit anti-MP-GAP antibody (1:250, self-made), Duolink *In Situ* PLA Probe Anti-Mouse MINUS (DUO92004, Sigma-Aldrich), Duolink *In Situ* PLA Probe Anti-Rabbit PLUS (DUO92002, Sigma-Aldrich), and Duolink *In Situ* Detection Reagents Orange (DUO92007, Sigma-Aldrich). Cells were seeded on 12 mm poly-L-lysine (Y1251-100G, Sigma) coated glass coverslips (P231.1, Carl Roth), synchronized with 0.5 mM thymidine (T1895, Sigma) for 24h, released and fixed 8-9 h later with 100% ice-cold methanol. Cells were blocked with provided blocking solution, incubated with primary antibodies against Aurora A and MP-GAP for 2 h at RT and washed with Wash Buffer (0.01 M Tris, 0.15 M NaCl and 0.05% Tween20, pH 7.4.) The two PLA probes were added and incubated for 1 h at 37°C. The probes were ligated together by adding ligation solution for 30 min at 37°C. Amplification buffer and polymerase were added to amplify positive signal for 1.5 h at 37°C, cells were washed with two wash buffers (Buffer A pH 7.4: 0.01M Tris, 0.15M NaCl, 0.05% Tween-29; Buffer B pH 7.5: 0.2M Tris, 0.1M NaCl), and co-stained with Hoechst 33258 (861405; Sigma-Aldrich). As a negative control same protocol was followed, however just one primary antibody against Aurora A or MP-GAP was used.

### Microscopy and image acquisition

For live-cell imaging cells were plated in ibidiTreat chambers on day one and incubated in medium containing 100 nM SiR-DNA (251SC007, Tebu Bio) for 60 min prior to image acquisition. Imaging was performed at 37°C in a heated chamber and cells were maintained in CO_2_-independent medium (18045054, Thermo Fischer) complemented with 10% FBS, 1% P/S, 0.2 mg/ml tetracycline, 2 mM L-Glutamine. MK-5108 and AZD treatments in HCT-116 during live-cell images were acquired on a Nikon eclipse Ti spinning disk confocal microscope. The Nikon was controlled by NIS Elements 4.51 and was equipped with a 100x 1.45-NA Plan-Apochromat oil immersion objective, a 488-, a 640-nm laser line, and an Andor DU-888 X-11056 camera (1,024 x 1,024 pixels). Other live-cell images (Fig. 6) were acquired on confocal DIVE-FALCON laser scanning confocal microscope (Leica Microsystems) controlled by the Leica LAS X Life Science software using the HCX PL APO CS 63.0×/1.30 NA glycerol immersion objective and 488 nm and 633 nm lasers. Immunofluorescence images were acquired on a Zeiss AxioImager.Z1 ApoTome microscope, which was controlled by AxioRel 4.6 software. The Zeiss AxioImager.Z1 was equipped with a 63x 1.4-NA Plan-Apochromat oil immersion objective (Zeiss) and AxioCamMR camera.

### Microscopy image analyses and quantifications

All fluorescence intensity measurements were performed using Fiji (ImageJ, Version 1.5) software. For anillin fluorescence intensity measurements a 6 pixel-wide linescan was drawn around the cell periphery from pole to pole (Fig. 1B). Additionally, a box adjacent to the cell pole was drawn to measure the cytoplasmic fluorescence intensity (CI). The fluorescence intensity profiles and mean cytoplasmic values were imported into KNIME (https://www.knime.com) for each cell and further processed. From all peripheral fluorescence intensity values the mean cytoplasmic values were subtracted. For metaphase cells, the mean fluorescence intensity over the whole cortex for each cell was calculated. For anaphase cells the mean polar intensity (0-10%, 40-60% and 90-100%) and equatorial intensity (20-30%, 70-80%) of total linescan length were calculated. To calculate the mean equatorial zone width running averages were calculated to smooth the fluorescence intensity curve and the zone width was calculated at half maximum fluorescence intensity for each zone of each cell at the cell equator.

MP-GAP cortical fluorescence intensity was measured by drawing two perpendicular 10 pixel-wide linescans across the cell periphery. One linescan was positioned at the cell equator and one from pole-to-pole (Fig. 4F). After plotting the fluorescence intensity data, the cortex peaks of MP-GAP could be assigned. The cytoplasmic MP-GAP intensities were calculated for each cell 5 pixel away from the two polar cortex peaks by averaging the measured intensities of the following 4 pixels. To normalize fluorescence intensities the measured cortex peak intensities were divided by the mean cytoplasmic intensity of each cell. For metaphase cells, the MP-GAP cortex factor was calculated by averaging the normalized intensities of the equatorial and polar linescan.

Spindle midzone length MK-5108 and AZD treatment was determined by measuring the distance between the two segregating chromosome masses at 6 min after anaphase onset in living HCT-116 cells (Fig. S1A) or during anaphase in fixed HeLa cells (Fig. S1E).

Cell elongation was quantified by measuring the pole-to-pole distance for each cell during anaphase over time (Fig. S2C). The measured distance of each timepoint was normalized to the distance at anaphase onset of the specific cell. Similarly, furrow ingression was analyzed by measuring the equatorial cell width for each cell during anaphase over time until complete furrow ingression (Fig. 3E). The measured equatorial cell width of each timepoint was normalized to the distance of the specific cell at anaphase onset.

### 6xHis-GAP and GST-MP-GAP fragments expression and purification

All 6xHis-tagged and GST-tagged proteins were expressed in 1L *E. coli* BL21(DE3) cultures. Protein expression was induced at an OD_600_ of 0.6-0.8 for 14h at 18°C with 0.1mM IPTG. Cells were harvested, bacterial pellets were washed once with cold 1x PBS and snap frozen in liquid N_2_. For the purification of 6xHis-tagged GAP domain (pEZ72 (Zanin et al., 2013)), bacterial pellets were thawn on ice and resuspended in lysis buffer (20 mM Tris-HCL pH 7.4, 500 mM NaCl, 0.2% Triton-X-100, 1 mM DTT (Roth, 6908.3), 1 mM PMSF (Sigma, p7626-1G), cOmplete protease inhibitor cocktail (Roche, 11836153001). Cells were lysed by sonification and clear lysates were incubated with Ni-NTA agarose beads (Qiagen, 30210) in binding buffer (50mM Tris-HCL, 500mM NaCl, 50mM Imidazol (Roth, 3899.1), 10% Glycerol) at 4°C for 1.5 hr. Lysates and beads were transferred into purification column, washed 3x 5min with 5x volumes binding buffer and proteins were eluted with elution buffer (50 mM Tris-HCL pH 7.4, 500 mM NaCl, 500 mM Imidazol, 10% Glycerol). Purified 6xHis-tagged proteins were dialysed at 4°C over-night with exchange buffer (10 mM Tris pH 8.0, 150 mM NaCl, 1 mM EDTA) using Slide-A-Lyzer^TM^ Dialysis Casette (ThermoFischer, 66380) following the manufactures instructions.

For purification of GST-tagged proteins, bacterial pellets were thawn on ice and resuspended in lysis buffer (10 mM Tris pH 8.0, 150 mM NaCl, 1 mM EDTA, 0.05% NP-40, 5 mM DTT, 1 mM PMSF, cOmplete protease inhibitor cocktail). Cells were lysed by sonification and clear lysates were incubated with glutathione Sepharose beads (Sigma, GE17-0756-01) at 4°C for 1.5 hr. Afterwards, beads were washed 3x 10min with lysis buffer and stored at 4°C.

### Pull down assay

For pull down assays 1 µg purified 6xHis-GAP domain was mixed with 2 µg purified Sepharose beads-bound GST-MP-GAP fragments in a total of 300 ml binding buffer (10 mM Tris pH 8.0, 150 mM NaCl, 1 mM EDTA, 10 mM Imidazol, 4 mM b-mercaptoethanol). Samples were rotated at 4°C for 1hr. Beads were washed 3×10min in binding buffer and resuspended in equal volumes of 2x sample buffer (0.1 M Tris, 4% SDS, 1.4% b-mercaptoethanol, 20% glycerol, 1% Bromphenol blue). Samples were cooked at 95°C for 5 min, run on a SDS-Page and stained with Coomassie and immunoblotted with anti-6xHis antibody (Invitrogene, MA1-135) to detect the bound 6xHis-GAP domain.

### Quantification and statistical analysis

The software R was used for statistical analyses (R Development Core Team, 2015) including the add-in package multcomp (Hothorn et al., 2008). First, data were examined for normal distribution using Shapiro-Wilk test. Datasets with two conditions with normal distributed data were analyzed by Student’s t-test and non-normally distributed data with Kruskal-Wallis Rank sum test. Datasets containing multiple conditions (>2) with non-equal distribution were analyzed by Kruskal-Wallis Rank sum test followed by multiple comparison post-hoc test. The corresponding statistical tests are indicated in the figure legends.

## Acknowledgments

Microscopy was performed either at the center for advanced light microscopy (CALM) at the LMU or at the optical imaging center Erlangen (OICE) at the FAU. We thank Helmut Brandstätter and Renato Frischknecht for access and introduction to the Apotom microscope. We are grateful to Elke Bachmann and Anna Lochner for their help with cloning the phosphomimetic MP-GAP variant. For critical comments on the manuscript we thank Shirin Bahmanyar and Mikhail Lebedev. The Deutsche Forschungsgemeinschaft supported E. Zanin (ZA619/3), and T. Mikeladze-Dvali (MI1867/1-3). The Leica SP8 DIVE-FALCON was supported by the DFG (INST 90/1074-1 FUGG for Benedikt Kost). S. Mangal was a member of International Max Planck Research School for Molecular Life Sciences and thanks the program for support. For sharing the HCT-116 NG-anillin cell line prior to publication we are grateful to Alisa Piekny.

---

**Figure S1.**
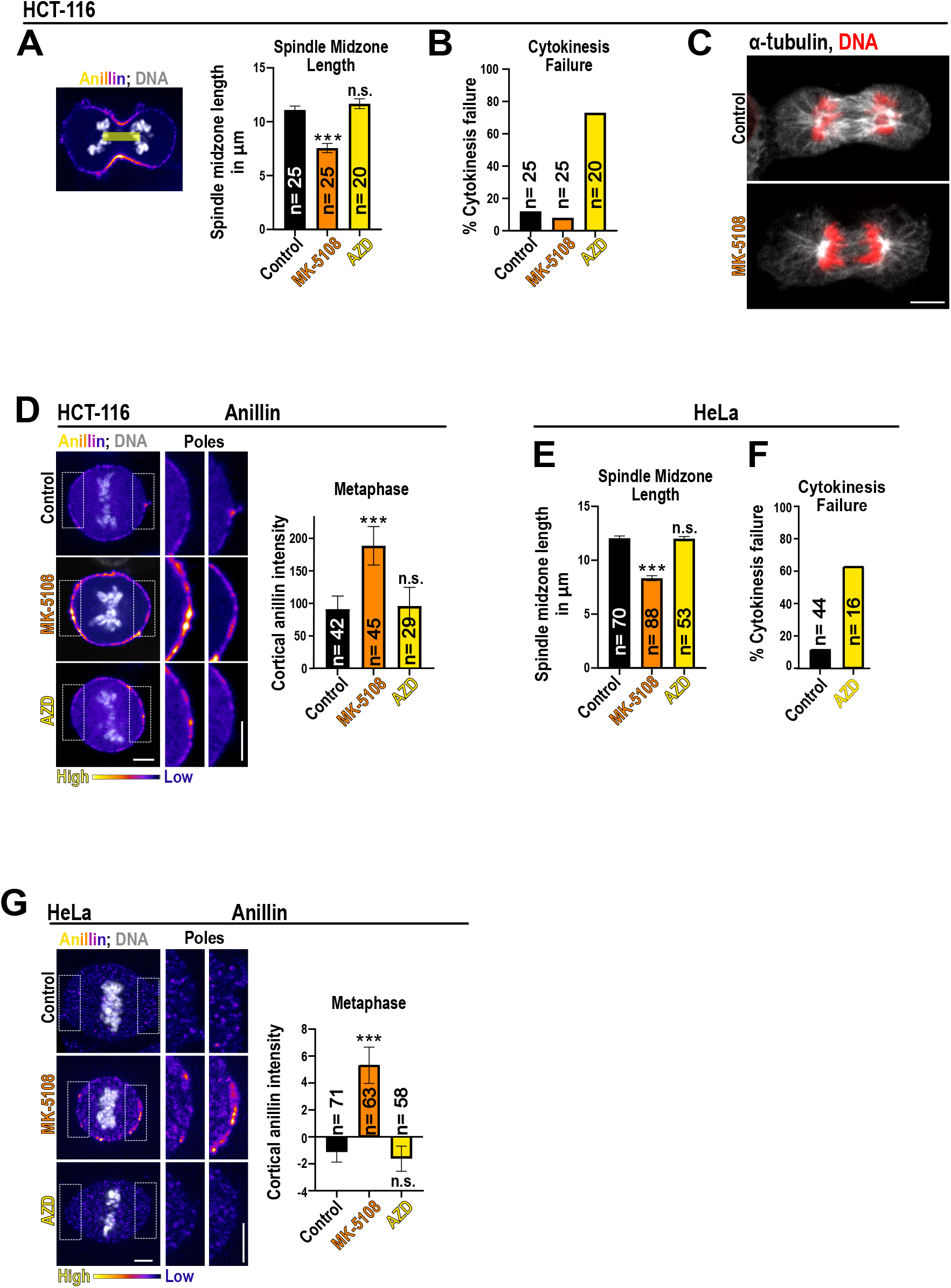
Addition of MK-5108 but not AZD increases cortical anillin levels in HCT-116 and HeLa cells during metaphase. A) Spindle midzone length was measured 6 min after anaphase onset in NG-Anillin expressing HCT-116 cells treated with DMSO (control), MK-5108 or AZD. B) Percentage of cytokinesis failure in NG-Anillin HCT-116 cells determined by live-cell imaging after treatment with the indicated inhibitors. C) Immunofluorescent images of HCT-116 cells treated with DMSO (control) or MK-5108 stained for α-tubulin (grey) and DNA (red). D) Shown are time-lapse fluorescent images of HCT-116 cells expressing NG-Anillin (pseudocolour) labeled for DNA (SiR-DNA, grey) in metaphase. HCT-116 cells were treated with the MK-5108, AZD, or DMSO (control). Insets show a magnification of the poles. Plotted is the mean normalized fluorescence intensity of anillin at the cortex during metaphase. E) Spindle midzone length of fixed HeLa cells in anaphase treated with DMSO, MK-5108 or AZD. F) Percentage of cytokinesis failure of HeLa cells determined by live-cell imaging after treatment with or without AZD. For all n = number of cells, and ≥2 independent experiments were performed for each condition. Error bars are SEM and p values were calculated with the Kruskal-Wallis test. Scale bars are 5 µm.

**Figure S2.**
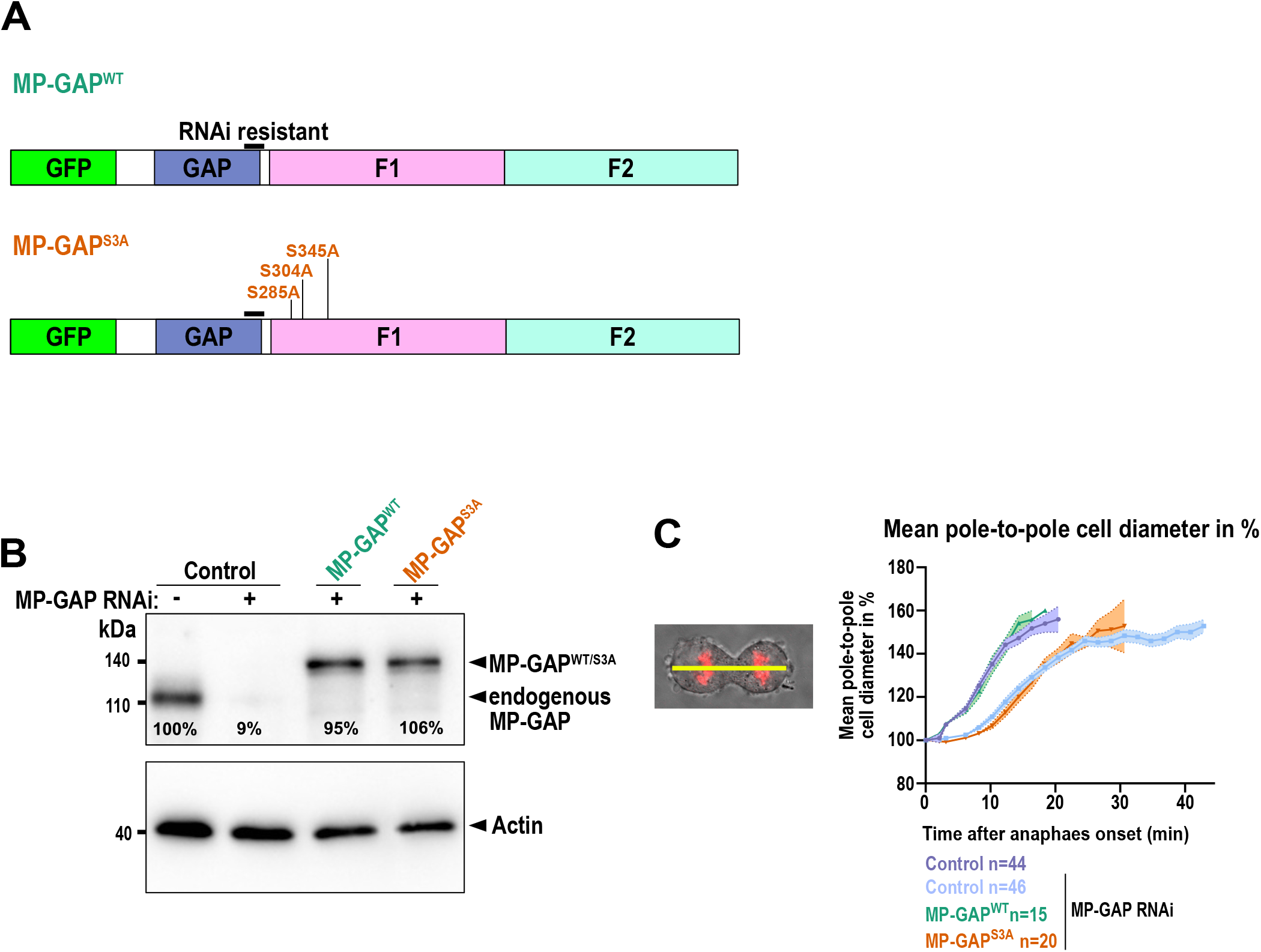
MP-GAP is required for cell elongation during anaphase. A) Scheme of the GFP-tagged RNAi resistant MP-GAP transgenes used. B) Immunoblot of control (without transgene) and GFP-tagged MP-GAP^WT^ and MP-GAP^S3A^ expressing HeLa cells for the indicated RNAi conditions probed with anti-MP-GAP and anti-actin antibody. Mean levels of endogenous MP-GAP, GFP-MP-GAP^WT^ or GFP-MP-GAP^S3A^ after MP-GAP RNAi relative to endogenous MP-GAP after control RNAi are shown for 3 independent extracts. C) Plotted is the mean pole-to-pole cell length over time for the indicated conditions.

**Figure S3.**
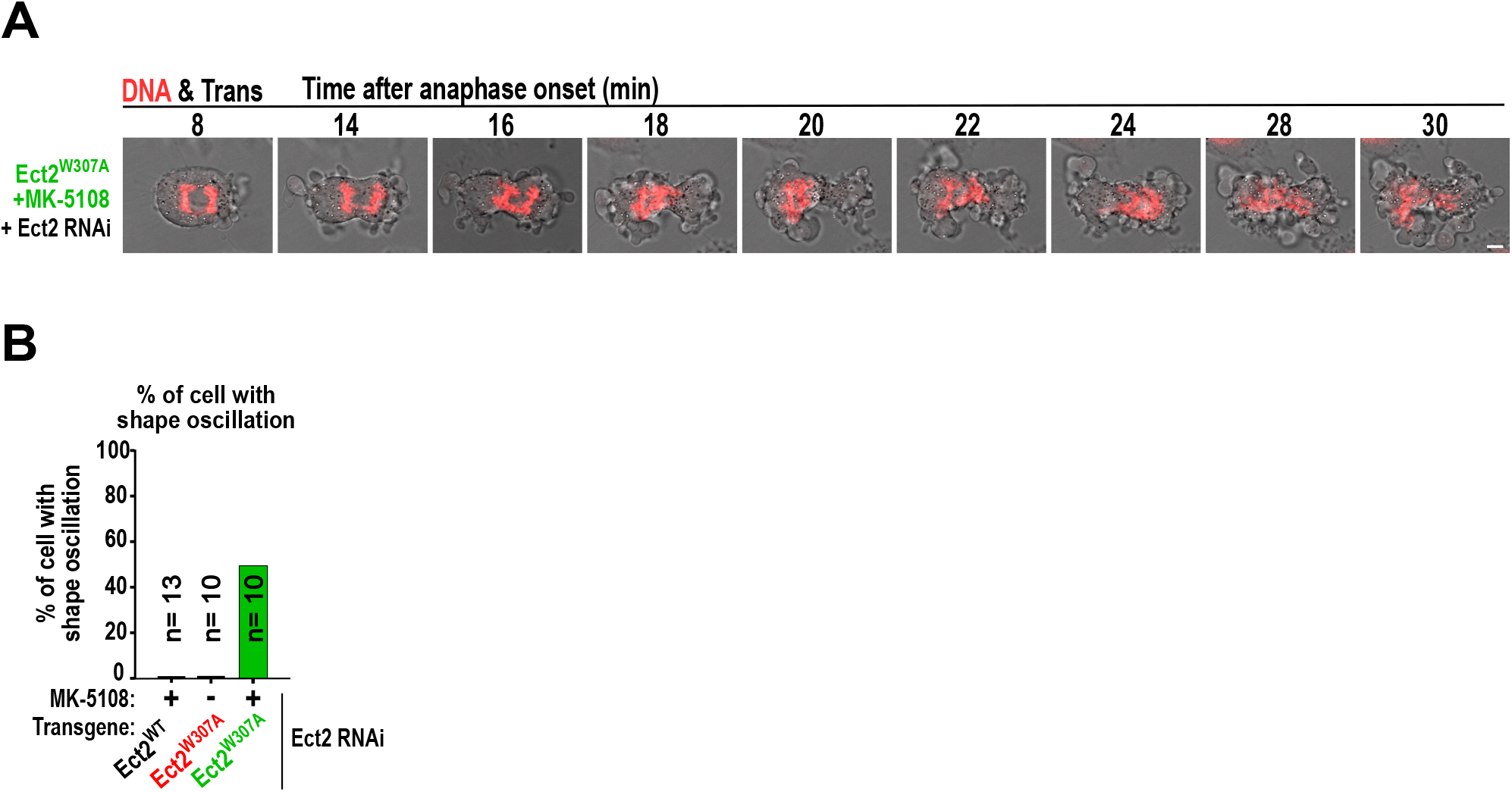
A) Merged transmission and DNA (SiR-DNA, red) time-lapse images of HeLa cell expressing Ect2^W307A^ and treated with MK-5108 around anaphase onset. Scale bar is 5 µm. B) Graph displays the percentage of HeLa cells with shape oscillation for the different treatment conditions. For all n = number of cells.

## Notes

### Competing Interest Statement

The authors have declared no competing interest.

